# A simplified *MiMe* for producing clonal gametes to enable synthetic apomixis in allotetraploid *Brassica napus*

**DOI:** 10.1101/2025.10.22.684060

**Authors:** Miaowei Geng, Lei Chu, Zilin Guo, Yashi Zhang, Chao Yang

## Abstract

Synthetic apomixis has been established in *Arabidopsis* and rice by combining *MiMe* that produces clonal unreduced gametes with the factors inducing parthenogenesis or haploids. The *MiMe* requires the simultaneous inactivation of three genes, which imposes challenges for its application in polyploids harbouring multiple copies for each gene. Here, we demonstrate that mutating two genes in the allotetraploid oil crop *Brassica napus* effectively converts meiosis into mitosis-like division, producing over 95% viable clonal gametes and thus establishing a simplified *MiMe* system in polyploids.

## Main text

Clonal reproduction through seeds (apomixis) in crops has great potential of application in agriculture, because it would allow indefinite self-propagation of any elite variety bearing genomic heterozygosity, in particular F1 hybrids. However, no major seed crop species has been shown to be able to reproduce naturally through apomixis. Recently, an engineered synthetic apomixis has been established in hybrid rice through combining the *MiMe* (mitosis instead of meiosis) that results in the production of clonal unreduced gametes, with either the ectopic expression of transcription factors specifically in egg cells triggering parthenogenesis, e.g., *BABY BOOM* (*BBM1* or *BBM4*) and *PARTHENOGENESIS* (*PAR*) or mutation of genes of haploid induction, e.g., *MATRILINEAL* (*MTL*) (Wang *et al*, 2019; Khanday *et al*, 2019; Song *et al*, 2024). The *MiMe* in Arabidopsis and rice is constructed by a combination of three recessive meiotic mutants, including the mutation of the cell-cycle regulator *omission of second division* (*osd1*) that produces unreduced gametes by skipping the second meiotic division, a second mutation that abolishes homologous pairing and recombination by preventing the DNA double strand break (DSB) formation (e.g., *spo11-1* in Arabidopsis, *pair1* in rice), and *rec8* that allows a precocious separation of sister chromatids at first meiotic division through rendering a bi-orientation of sister kinetochores (Wang, 2019). While *MiMe* is likely functionally conserved in a broad range of crops (Mieulet *et al*, 2016), the requirement of a simultaneous mutation of these three genes in one generation imposes extreme challenges for its wide application in engineering the synthetic apomixis of crops, especially of polyploids where each target gene has two or even more equivalent copies.

In this study, we aimed to establish the *MiMe* in an allotetraploid species *Brassica napus*. A phylogenetic analysis for *SPO11-1*, *REC8*, and *OSD1* reveals that they all have two homologs in *Brassica napus*, with one from A subgenome and another one from C subgenome (Figure S1a-c). To this end, we first sought to generate the *spo11-1*, *rec8*, and *osd1* mutants independently by CRISPR/Cas9 in the spring-type rapeseed cultivar *Westar*. Two independent homozygous mutant lines were identified from T1 generation for each target gene (Figure S2a-c). We further confirmed the knockout of *SPO11-1* or *REC8* by immunostaining using DMC1 or REC8 antibodies (Figure S2d,e). Similar to that in *Arabidopsis* and rice, mutations of *spo11-1* and *rec8* exhibited no effect on vegetative growth of *Brassica napus*, consistent with their meiosis-specific roles (Figure S3a). We also did not observe obvious growth defect in *osd1* mutants (Figure S4a).

During the reproductive stage, *spo11-1* mutants were largely infertile, whereas *rec8* mutation led to complete sterility, as shown by the reduced or abolished seed sets and pollen viability in *spo11-1* and *rec8* mutants, respectively (Figure 1a-d, Figure S3b). As expected, *osd1* mutants showed a wild-type level of fertility. However, differing from *osd1* in Arabidopsis, the sizes of pollen grains in *osd1* of *Brassica napus* appear normal (Figure S4b). Chromosome spreads and tetrad staining of *osd1* also revealed a comparable meiotic program to that in wildtype, without skipping the second meiotic division, indicating likely the functional redundancy of *OSD1* with other factors, such as its paralog *UVI4* (Figure S4c-e).

**Figure 1.**
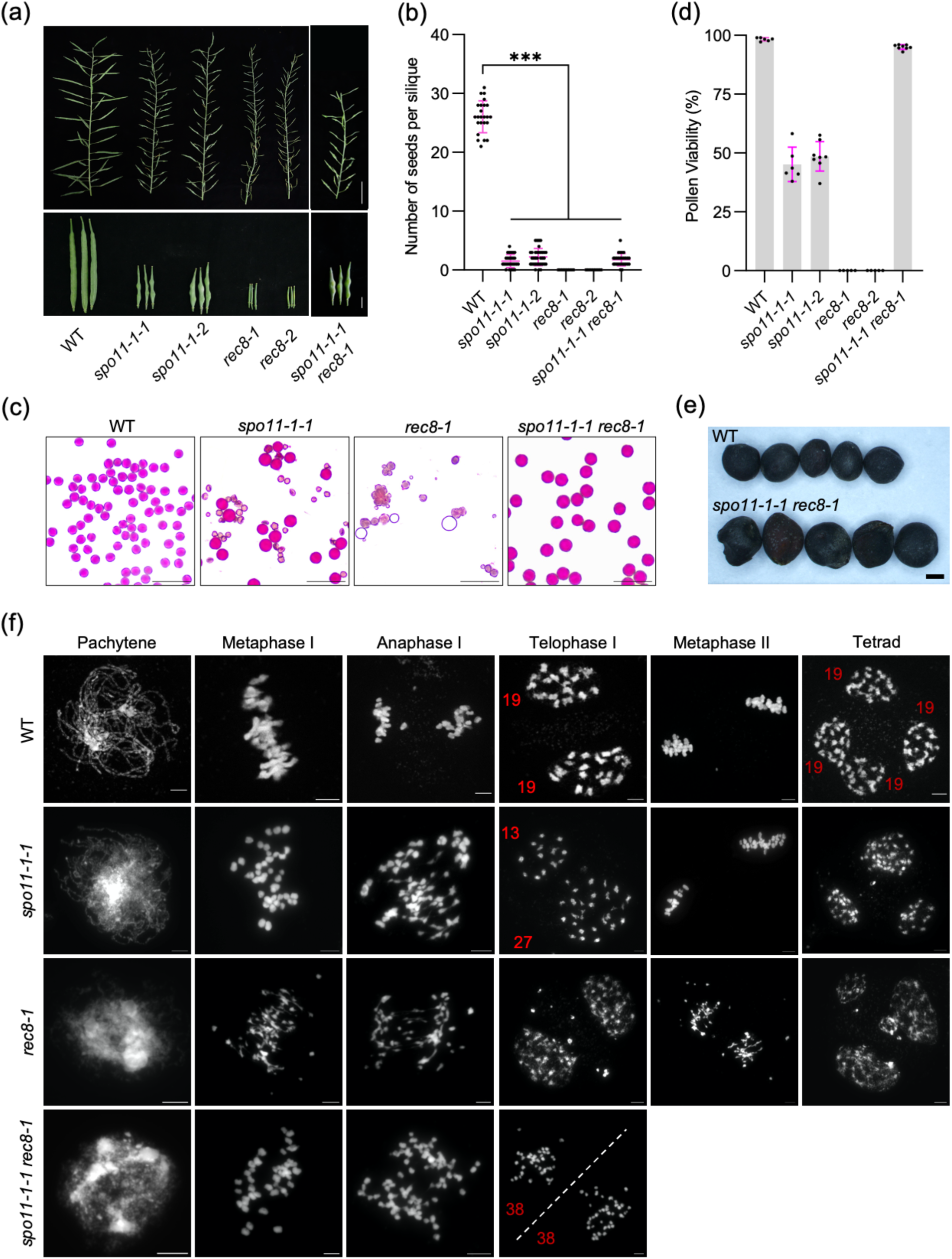
Phenotypic analysis of plant fertility and meiotic behaviour in WT, *spo11-1*, *rec8* and *spo11-1 rec8-1* mutants. (a) Main branches and siliques in WT, *spo11-1-1*, *spo11-1-2*, *rec8-1*, *rec8-2* and *spo11-1 rec8-1* mutant plants. The bars from top to bottom are 5 cm and 1 cm, respectively. (b) Statistical analysis for the number of seeds per silique in WT, *spo11-1-1*, *spo11-1-2*, *rec8-1*, *rec8-2* and *spo11-1 rec8-1* mutant plants. At least 24 siliques were counted for each genotype. (c) Pollen staining of WT, *spo11-1-1*, *rec8-1* and *spo11-1 rec8-1* mutant plants. Red colour indicates viable pollen grains and grey indicates aborted ones. Bars: 100µm. (d) Statistical analysis for the pollen viability of WT, *spo11-1-1*, *spo11-1-2*, *rec8-1*, *rec8-2* and *spo11-1 rec8-1* mutant plants. At least 6000 pollen grains were counted for each mutant. (e) Seeds of WT and *spo11-1 rec8-1* mutant. Bars: 1mm. (f) Chromosome spreads of male meiosis in WT, *spo11-1-1*, *rec8-1* and *spo11-1-1 rec8-1* mutants throughout meiosis. Bars: 5µm. Asterisks in (b and d) indicate significant difference (Tukey’s multiple comparison test, *p*<0.001).

Next, we crossed *spo11-1-1* and *rec8-1* and obtained *spo11-1 rec8* double mutants from F3 segregating populations. Unexpectedly, in contrast to the strongly defective pollen viability in *spo11-1* and *rec8*, a nearly wild-type level of pollen viability was observed in *spo11-1 rec8* double mutants (98.3% in WT vs 95.0% in *spo11-1 rec8*) (Figure 1c,d), which is markedly different from the drastic reduction in *Arabidopsis spo11-1 rec8* or rice *pair1 rec8* mutants (Mieulet *et al*, 2016; Chelysheva *et al*, 2005). The pollen grains generated in *spo11-1 rec8* of *Brassica napus* are similar in size, which, however, are larger than that in the wildtype, suggesting likely an increased ploidy (Figure 1c). This is also supported by the larger seeds produced by *spo11-1 rec8*, despite only a few per silique (Figure 1e). These results suggest a marked difference in meiotic progression between *Brassica napus spo11-1 rec8* and *Arabidopsis spo11-1 rec8* or rice *pair1 rec8*.

Therefore, we investigated the impact of *spo11-1* and *rec8* mutations on meiosis in *Brassica napus*. We first analyzed male meiotic products at tetrad stage using orcein staining. While the wildtype produces mostly balanced tetrads (98.8%), the mutation of *spo11-1* and *rec8* produce mainly defective tetrads, e.g., monads, triads, unequal tetrads, and polyads, indicating a major problem in meiosis (Figure S3c,d). In contrast, *spo11-1 rec8* double mutants produced mainly balanced dyads (98.0%), consistent with their production of viable and large pollens, suggesting an omission of the second meiotic division (Figure S3c,d).

We then performed chromosome spreads to analyze the chromosome behavior in male meiosis. In the wildtype, homologous chromosomes (homologs) paired and synapsed at pachytene, which were further condensed to form 19 bivalents aligning at the metaphase I plate (Figure 1f). For each bivalent, the homologs were linked by at least one crossover formed through DSB repair, and were segregated equally into opposite poles at anaphase I, thus reducing chromosomes by half. During second meiotic division sister chromatids separate from each other to produce balanced tetrads where each daughter cell contains 19 chromatids (Figure 1f).

In *spo11-1*, the homologs cannot pair and recombine due to the absence of DSBs, resulting in the presence of 38 univalents at metaphase I. At anaphase I, univalents were mostly pulled in random to one pole or sometimes sister chromatids showed a precocious separation, leading to the formation of defective tetrads after second meiotic division (Figure 1f, Figure S5a-c). In *rec8*, an asynaptic behavior was also seen, and chromosome fragments were prevalent at metaphase I, producing hardly any normal tetrads, confirming the importance of REC8 for chromosome pairing and DSB repair in *Brassica napus* (Figure 1f, Figure S5a).

In *spo11-1 rec8*, the chromosome fragments present in *rec8* were not observed, and instead, 38 univalents appeared at metaphase I plate as *in spo11-1*, corroborating the abolishment of DSB formation and recombination in the absence of SPO11-1 (Figure 1f). From anaphase I to telophase I, all sister chromatids showed a mitosis-like equal separation into two groups (38:38) in all observed cells (n=29), indicating a conserved role of REC8 in regulating the mono-orientation of sister kinetochores in *Brassica napus* as in *Arabidopsis* and rice (Figure 1f). We further confirmed the equal distribution of A and C subgenomes by FISH using a C subgenome-specific probe (Figure S5d). Unexpectedly, after the first meiotic division, we hardly observe any configuration of second meiotic division in *spo11-1 rec8*, suggesting the omission of meiosis II, thereby resulting in formation of clonal unreduced gametes (Figure 1f, Figure S3c,d). This mitotic-like division in *spo11-1 rec8* of *Brassica napus* mimics the *MiMe* in *Arabidopsis* (*spo11-1 rec8 osd1*) and rice (*pair1 rec8 osd1*).

Next, we asked whether female meiocytes also produce clonal gametes in *spo11-1 rec8* of *Brassica napus*. Because the female meiosis in plants is difficult to observe, we sought to verify this question indirectly by assessing the ploidy of the offspring of *spo11-1 rec8*. Indeed, all analyzed progenies (n=20) showed the phenotypes of late flowering and enlarged floral organs that are typical indicators of increased ploidy, and 76 chromosomes (octoploid, 8n) were presented in metaphase I cells (Figure S6a-c). This 8n ploidy was further confirmed by flow cytometry, suggesting that female meiocytes in *spo11-1 rec8* also turn meiosis into mitosis (Figure S6d). Furthermore, these octoploid progenies (called *spo11-1 rec8* (8n)) produce more enlarged pollen grains compared to that in the tetraploid *spo11-1* rec8, while having still a relatively high viability at ∼ 85% (Figure S6e,f). Consistently, cytological analyses revealed a mitotic-like meiotic program and high proportion of dyads (∼ 84.7%) in the octoploid *spo11-1 rec8* (8n), thus producing clonal unreduced gametes with 76 chromatids (8n) (Figure S4c,g). Notably, the reciprocal crossing experiments between *spo11-1 rec8* and wildtype demonstrated that the low seed set in *spo11-1 rec8* is unlikely caused by problems in female meiosis, but rather attributed to the failure of fertilization and/or early embryogenic defects induced potentially by the increased ploidy and high chromosome numbers (Figure S3e).

Here, in contrast to *Arabidopsi*s and rice, we demonstrate that the *MiMe* in *Brassica napus* requires only mutations of two genes, namely *REC8* plus one gene required for DSB formation (e.g., *SPO11-1*, *SPO11-2*, *MTOPVIB*, *PRD1*, *PRD2*, *PRD3*, or *DFO*). We further validated this approach in the background of F1 hybrids of two rapeseed cultivars (*Westar* x *J9707*) by targeting two gene combinations (*SPO11-1* or *MTOPVIB* with REC8) using CRISPR/Cas9, producing a ratio of ∼ 92.0% and 93.5% clonal gametes, respectively and 8n progenies that show a non-segregating growth (Figure S7a-e, Figure S8). The fix of heterozygosity in these 8n progenies was verified further by whole-genome resequencing (Figure S9).

This new *MiMe* for *Brassica napus* provides at least two benefits, i.e., elevating the efficiency of the construction of *MiMe* because it does not require to mutate *OSD1*, which in turn avoids the potential adverse effects on plant growth induced by the absence of OSD1 that has been shown to affect mitotic progression and cell fate in *Arabidopsis* (Iwata *et al*, 2011; Singh *et al*, 2015). The underlying reason for skipping second meiotic division without needing an *osd1* mutation in this *Brassica napus MiMe* remains unclear. This might be related to the (allo)polyploidy and/or high chromosome numbers, which requires further investigation. Considering the conserved functions of SPO11-1 and REC8 across different sexual eukaryotes, it is compelling to study the feasibility of this *MiMe* established in *Brassica napus* in other polyploid crops such as wheat. The wide application of this technology in *Brassica napus* and other crops requires the development of high-throughput transgenic approaches and efficient gene editing tools. In the future, combining this *MiMe* with the technique of parthenogenesis or haploid induction would allow clonal seed production in *Brassica napus*.

## Materials and methods

### Plant materials and growth condition

The spring-type *Brassica napus* cultivar *Westar* and its F1 hybrid with another spring-type variety *J9707* (*Westar* × *J9707*) were used as the wild-type reference in this study. The *spo11-1*, *rec8*, and *osd1* mutants were generated by CRISPR/Cas9 gene editing technique. The *spo11-1-1 rec8-1* double mutants were identified from the F3 populations of the cross of *spo11-1-1* with *rec8-1*. The rapeseed plants were grown in a greenhouse with a condition of 16 h light at 22°C and 8 h dark at 20°C.

### Construction of CRISPR/Cas9 vector

To generate *Brassica napus spo11-1* and *rec8* mutants, two sgRNAs targeting the conserved sequences of two exons for both *SPO11-1* and REC8 were designed using the CRISPR-P2.0 (http://crispr.hzau.edu.cn/CRISPR2) (Figure S1b,c). The sgRNAs were synthesized and inserted into the binary vector *PKSE401* that contains the guide RNA scaffold and Cas9 expression cassettes, using the golden gate assembly method (Xing *et al*, 2014). To construct the CRISPR/Cas9 vectors that simultaneously target *SPO11-1* and *REC8* in F1 hybrids (*Westar* x *J9707*), one of these two sgRNAs according to their editing efficiency were selected, and then integrated into one T-DNA vector (*PKSE401*). To construct the CRISPR/Cas9 vectors that simultaneously target *MTOPVIB* and *REC8* in F1 hybrids (*Westar* x *J9707*), a new sgRNA targeting *MTOPVIB* was designed and inserted into the PKSE401 vector harboring the sgRNA for *REC8* (Figure S5a). The primers for creating these vectors are listed in Supplemental table 1.

### Plant transformation and genotyping

The agrobacterium-mediated transformation of *Brassica napus* was carried out as previously described using hypocotyl as explants (Dai *et al*, 2020). To identify the mutations induced by CRISPR/Cas9, each copy of *SPO11-1* and *REC8* was amplified by PCR using the gene-specific primers. The PCR fragments were then sequenced by the gene-specific primers. The primers used are listed in Supplemental table 1.

### Pollen and tetrad staining

The viability staining of pollen was performed by dipping open flowers into Peterson staining solution on microscopic slides, which were then heated at 80°C for 10 mins to increase the contrast between dead and viable pollen grains (Peterson *et al*, 2010). For tetrad analysis, the flower buds of appropriate size were dissected, and isolated anthers were softened in 1 mol/L HCl at 60°C for 1 minute. Subsequently, anthers were rinsed in distilled water and squashed in orcein staining solution on slides for microscopic examination. Images were captured using a SOPTOP optical microscope ex30 equipped with a color camera.

### Chromosome spread

Chromosome spreading analysis was carried out as described previously with slight modifications (Chelysheva *et al*, 2010). Flowers at early flowering stage were collected and fixed in ethanol: acetic acid (3:1, v/v) fixative for 72h at 4℃, and then stored at −20℃. To prepare the chromosome spreads, the entire flower was washed in sodium citrate buffer (pH 4.5) for three times (10 minutes each) to remove the fixative. Flower buds of appropriate size (∼1-2mm) were selected. From each bud, one anther was taken and squashed in Carbol-fuchsin solution to deterime the meiotic stage under a light microscope. Next, the remaining 5 anthers were placed in an enzyme solution (3% cellulose, 3% mazerozyme, and 5% snailase in 50 mM citrate buffer pH 4.5) to digest for 50 minutes. Subsequently, single digested anthers were transferred onto a microscopic slide, 15 µl of 60% ice-cold acetic acid was added, and anthers were then macerated into a homogenous mixture using a bent needle. An additional 15 µl of 60% ice-cold acetic acid were added and the liquid mixture was spread immediately by circular stirring on a heating block at 45°C until the liquid was largely evaporated. The slide was rinsed once with ice-cold fixative (3:1 v/v ethanol: acetic acid), drained shortly on the heat block, and dried at room temperature for 1 hour. The slides were then stained with DAPI for observing meiotic chromosome behaviors. Images were captured using a SOPTOP fluorescence microscope RX50 equipped with a monochrome camera.

### Flow cytometry

To verify the cell ploidy by flow cytometry, a piece of fresh leaf (∼ 100mg) was chopped into homogenate with a sharp blade in a small petri dish with 200 µl of precooled nuclei extraction buffer containing DAPI. After 1-2 mins incubation, 1.5 ml water was added into the petri dish and then the extracts were filtered through a 50 μm filter. The DAPI stained nuclei extracts were analysed by CyFlow Ploidy Analyser (Sysmex).

### Whole-genome resequencing and genotype calling

Genomic DNA was extracted from young leaves using the CTAB method. Sequencing libraries were constructed using the VAHTS™ Universal DNA Library Prep Kit for Illumina® V3 (Vazyme Biotech) according to the manufacturer’s manual. Paired-end sequencing was performed on the Illumina Nova 6000 platform, with a read length of 150 bp. The raw sequencing reads were filtered by the fastp software to generated high-quality clean reads. Then, the clean reads from individuals were aligned to the reference genome (*Westar*, http://cbi.hzau.edu.cn) using the bwa mem with default parameters. Only the uniquely mapped reads with both ends located within a genomic region < 1 Kb were kept for subsequent analysis. The alignment results in SAM format were converted to the BAM format and then sorted using SAMtools. Duplicated reads caused by PCR in the process of library construction were identified and removed using Picard package. Variant (including SNP and InDel) calling for all accessions was performed using the HaplotypeCaller module in GATK software. To obtain high-quality variants, raw variants were further filtered with the following criteria: (a) bi-allelic, homozygous and polymorphic between parents; (b) the sequencing depth should be larger than 3; (c) the percentage of missing genotype is less than 50%. Finally, we obtained 1433268 high-quality markers. A sliding window-based method was used to construct the genome-wide genotype landscape for all samples using a 200 Kb window size and 100 Kb window step.

## Accession numbers

The sequence data can be found in the Brassica napus multi-omics information resource database BnIR under the following accession numbers*: BnaA01.SPO11-1* ( BnaA01G0362000ZS), *BnaC01.SPO11-1* (BnaC01G0452000ZS), *BnaA01.MTOPVIB* (BnaA01G0272200ZS), *BnaC01.MTOPVIB* (BnaC01G0332400ZS), *BnaA10.REC8* (BnaA10G0274700ZS), *BnaC09.REC8* (BnaC09G0592200ZS), *BnaA09.OSD1* (BnaA09G0532300ZS), and *BnaC08.OSD1* (BnaC08G0378700ZS).

## Acknowledgements

This work was supported by the National Key Research and Development Program of China (2023YFF1000700), National Natural Science Foundation of China (32370360, 32170354), Science and Technology Innovation 2030 Major Project of China (2022ZD04010), Hubei Key Research and Development Program (2023BBB173), and Fundamental Research Funds for the Central Universities (2662025ZKPY004).

## Author contributions

C.Y. conceived this research. M.G., L.C., Z.G., and Y.Z. performed the experiments and analyzed the data. C.Y. and M.G. wrote the manuscript. All authors commented, discussed and provided input on the paper.

## Data availability statement

The data that supports the findings of this study are available in the supplementary material of this article.

## Supplemental figures

**Figure S1.**
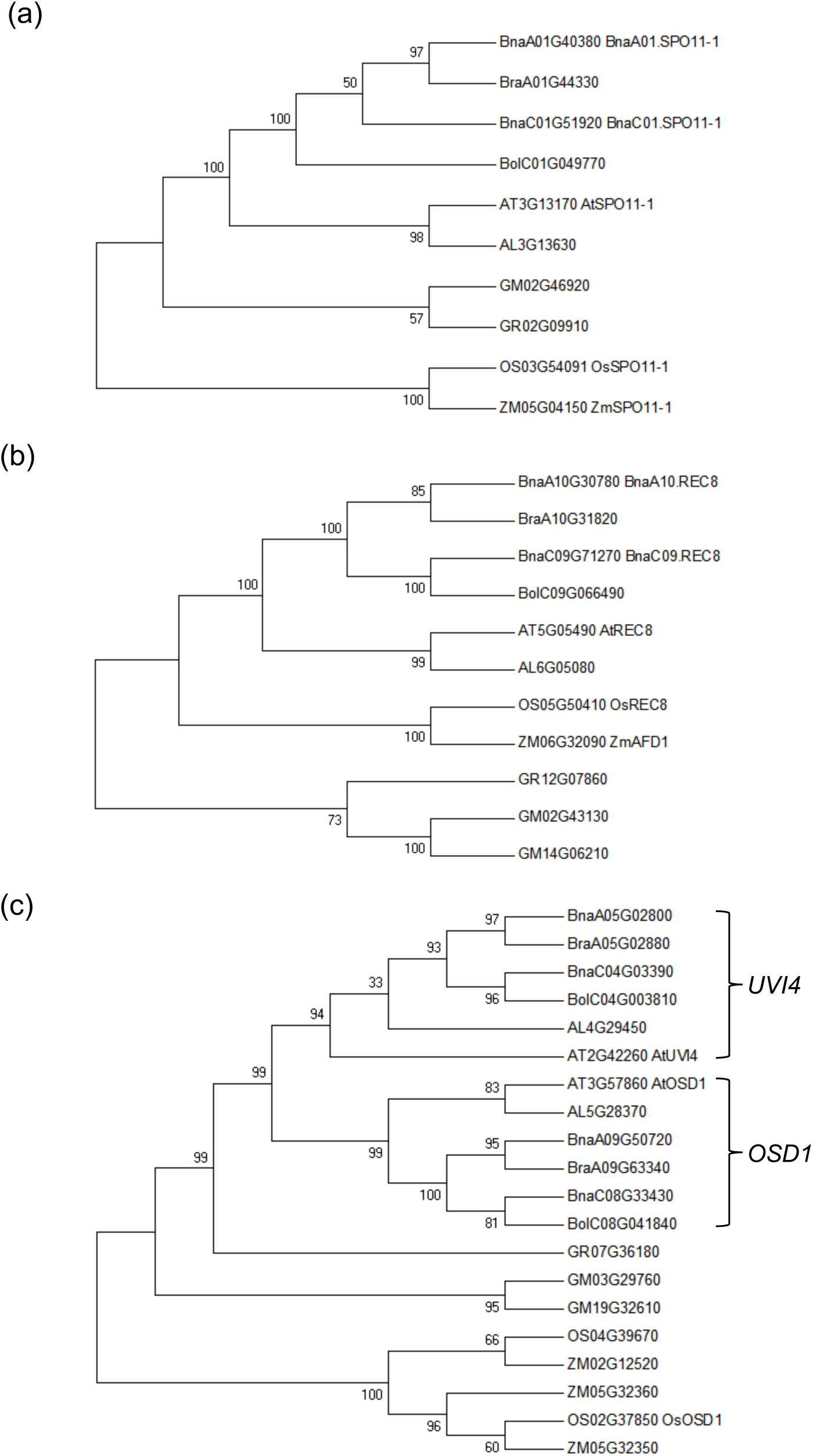
Phylogenetic analyses for SPO11-1, REC8 and OSD1 homologues. Unrooted phylogenetic tree was built based on the full-length protein alignment of SPO11-1 (a), REC8 (b), and OSD1/UVI4 (c) family proteins using the maximum likelihood method. Proteins from *Arabidopsis thaliana* (At), *Arabidopsis lyrata* (Al), *Brassica napus* (Bna), *Brassica rapa* (Bra), *Brassica oleracea* (Bol), *Glycine max* (Gm), Oryza sativa (Os), and *Gossypium raimondii* (GR).

**Figure S2.**
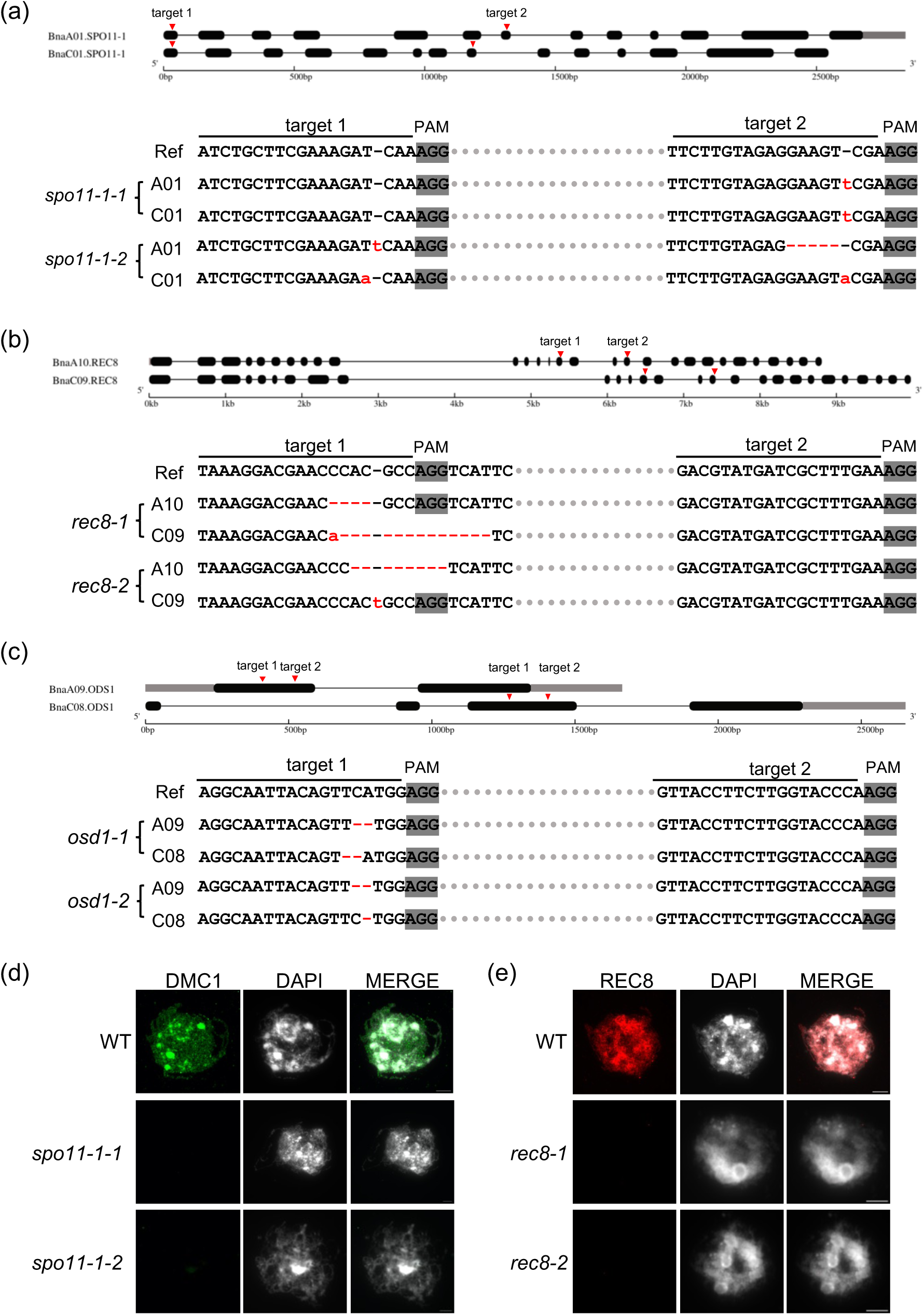
Generation of *spo11-1*, *rec8*, and *osd1* mutants in *Brassica napus.* (a) Generation and identification of two *spo11-1* mutant lines by CRISPR-Cas9. (b) Generation and identification of two *rec8* mutant lines by CRISPR-Cas9. (b) Generation and identification of two *osd1* mutant lines by CRISPR-Cas9. Red arrowheads in (a-c) indicate the targeting regions of sgRNAs. (d) Immunostaining of DMC1 in wildtype, *spo11-1-1*, and *spo11-1-2* mutants. Bars: 5µm. (e) Immunostaining of REC8 in wildtype, *rec8-1*, and *rec8-2* mutants. Bars: 5µm.

**Figure S3.**
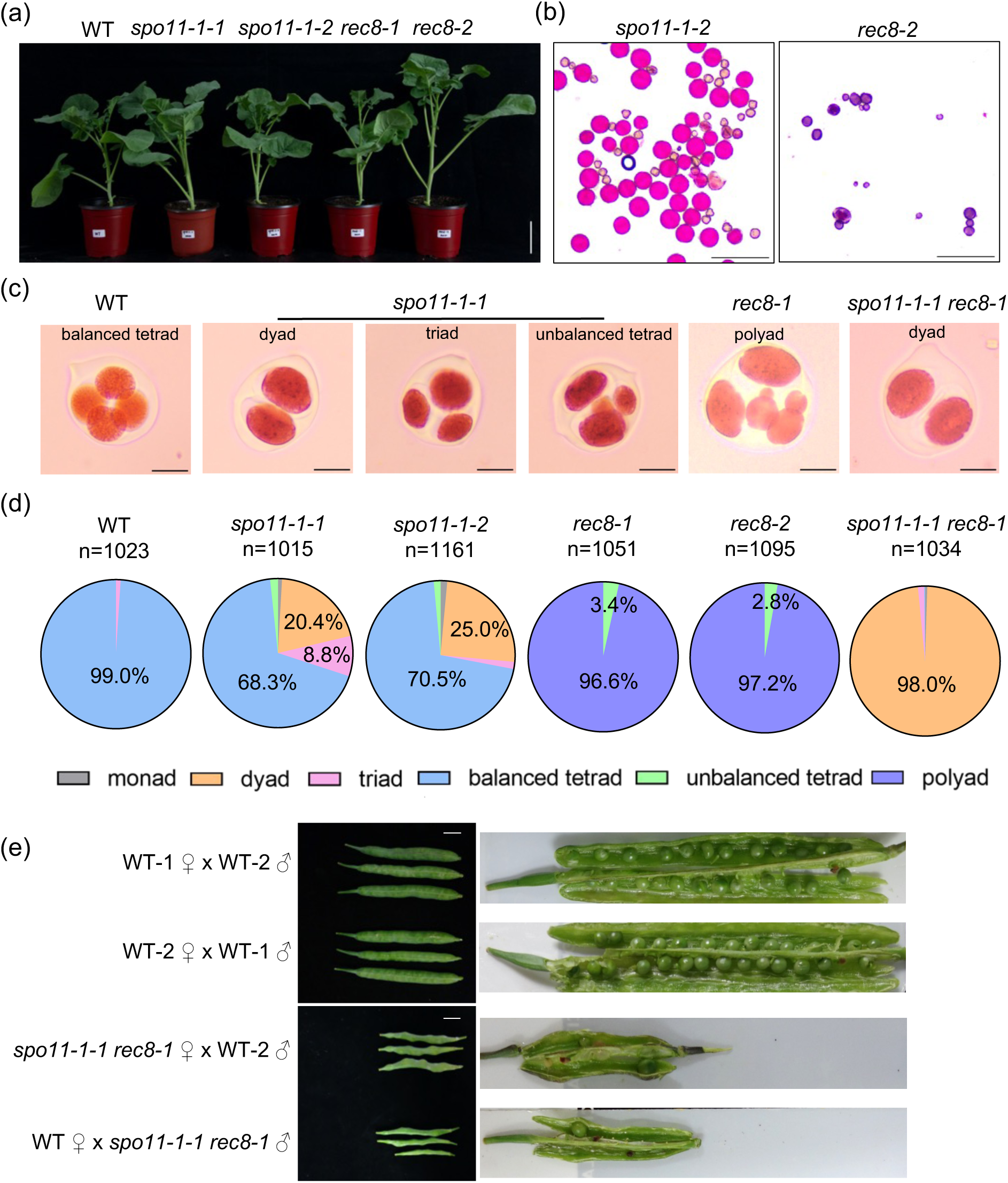
Phenotypic analyses of plant growth and male meiotic products of WT, *spo11-1*, *rec8* and *spo11-1 rec8-1* mutants. (a) The vegetative growth of WT, *spo11-1-1*, *spo11-1-2*, *rec8-1*, *rec8-2*, and *spo11-1 rec8-1* mutants. Bar: 10cm. (b) Pollen staining of *spo11-1-2* and *rec8-2* mutant plants. Red pollen grains are fertile and grey pollen grains are aborted. Bars: 100µm. (c) Representative images of male meiotic products at tetrad stage in WT, *spo11-1*, *rec8-1*, and *spo11-1 rec8-1* mutants. Bars: 5µm. (d) Pie charts showing the proportion of different meiotic products including monad, dyad, triad, balanced tetrad, unbalanced tetrad, and polyad in WT, *spo11-1-1*, *spo11-1-2*, *rec8-1*, *rec8-2*, and *spo11-1-1 rec8-1* mutant plants. (e) Siliques from the reciprocal crossings between *spo11-1-1 rec8-1* and WT. Bars: 1cm.

**Figure S4.**
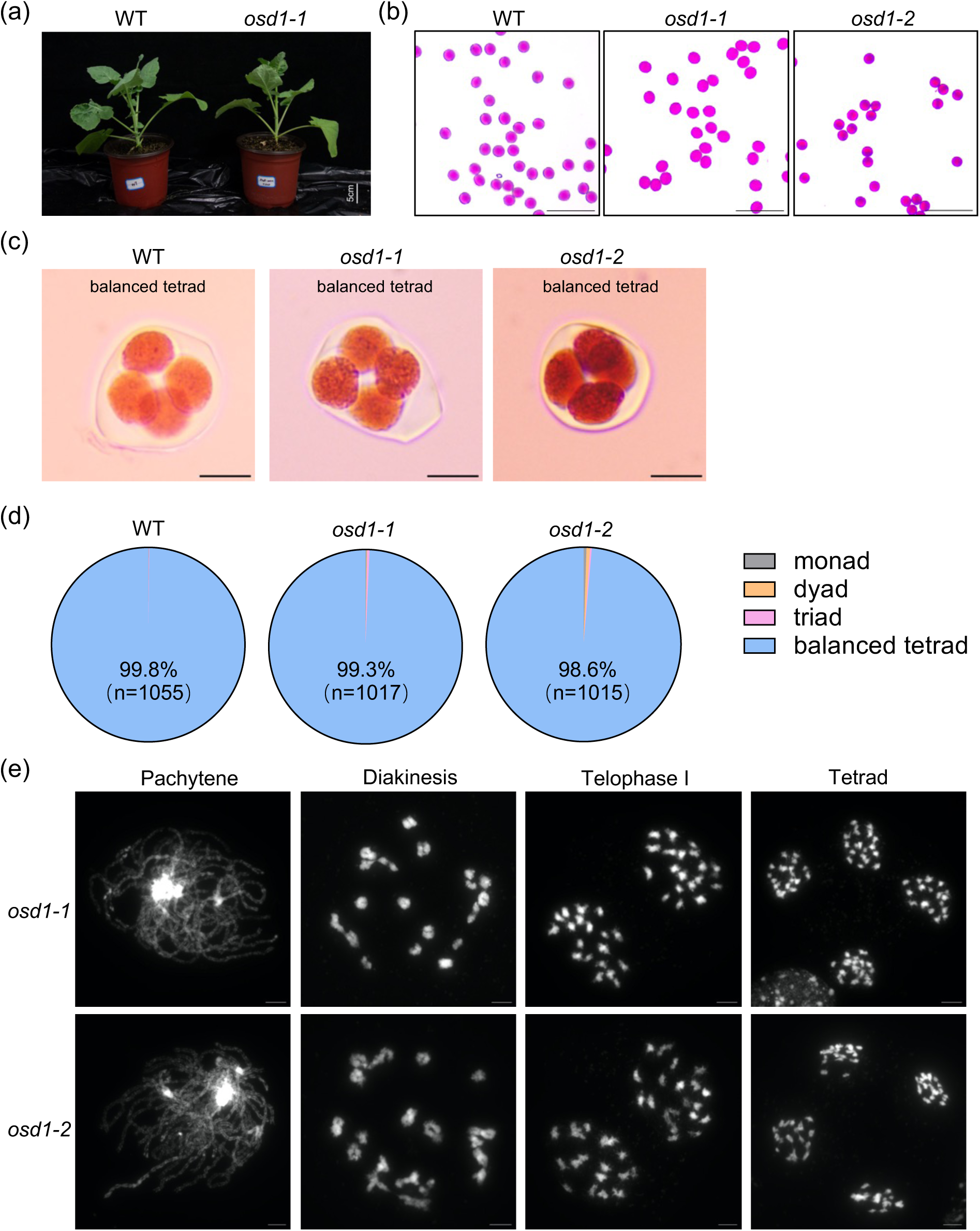
Phenotypic analyses of plant growth and male meiotic products of WT, *osd1-1*, and *osd1-2* mutants. (a) The vegetative growth of WT and *osd1-1* mutant. Bar: 10cm. (b) Pollen staining of *spo11-1-2* and *rec8-2* mutant plants. Red pollen grains are fertile and grey pollen grains are aborted. Bars: 100µm. (c) Representative images of male meiotic products at tetrad stage in WT, *osd1-1*, and *osd1-2* mutants. Bars: 5µm. (d) Pie charts showing the proportion of different meiotic products including monad, dyad, triad, and balanced tetrad in WT, *osd1-1*, and *osd1-2* mutant plants. (e) Chromosome spread analysis of male meiosis in *osd1-1* and *osd1-2* mutants. Bars: 5µm.

**Figure S5.**
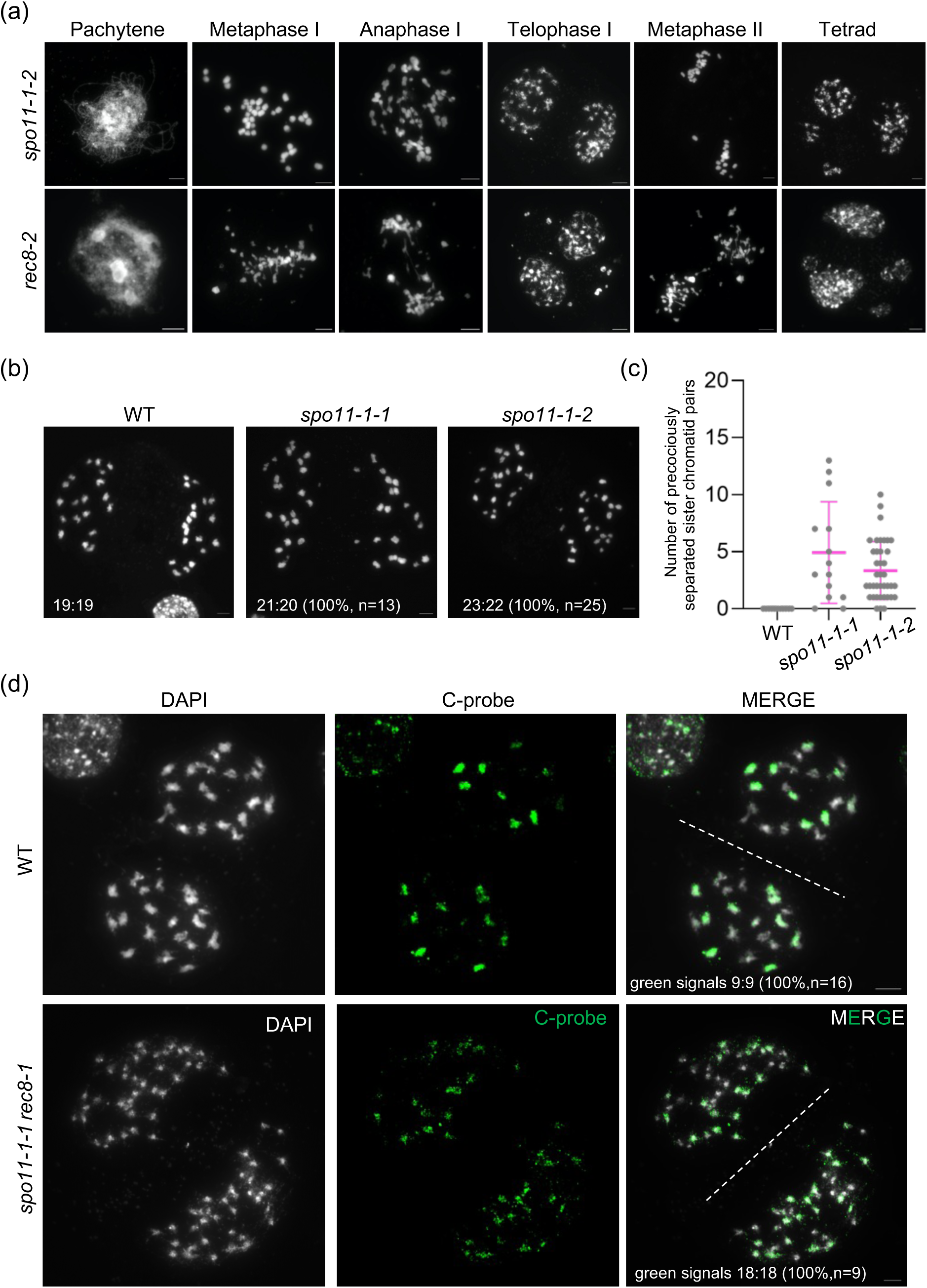
Meiotic behavior analysis in WT, *spo11-1-1*, *spo11-1-2*, *rec8-2*, and *spo11-1-1 rec8-1* mutant plants. **(a)** Chromosome spreads of male meiosis in *spo11-1-2* and *rec8-2* mutants throughout meiosis. Bars: 5µm. (b) Analysis of chromosome distribution at telophase I in male meiosis of WT, *spo11-1-1*, and *spo11-1-2* mutants. No balanced segregation was observed in *spo11-1-1* and *spo11-1-2* mutants. Bars: 5µm. (c) Quantification of the precociously separated sister chromatid pairs during meiosis I in WT, *spo11-1-1*, and *spo11-1-2* mutants according to (b). (d) FISH analysis of the distribution of A and C subgenomes in WT and *spo11-1-1 rec8-1* mutants at telophase I of telophase I-like stage. The green signals indicate the chromosomes from C subgenome.

**Figure S6.**
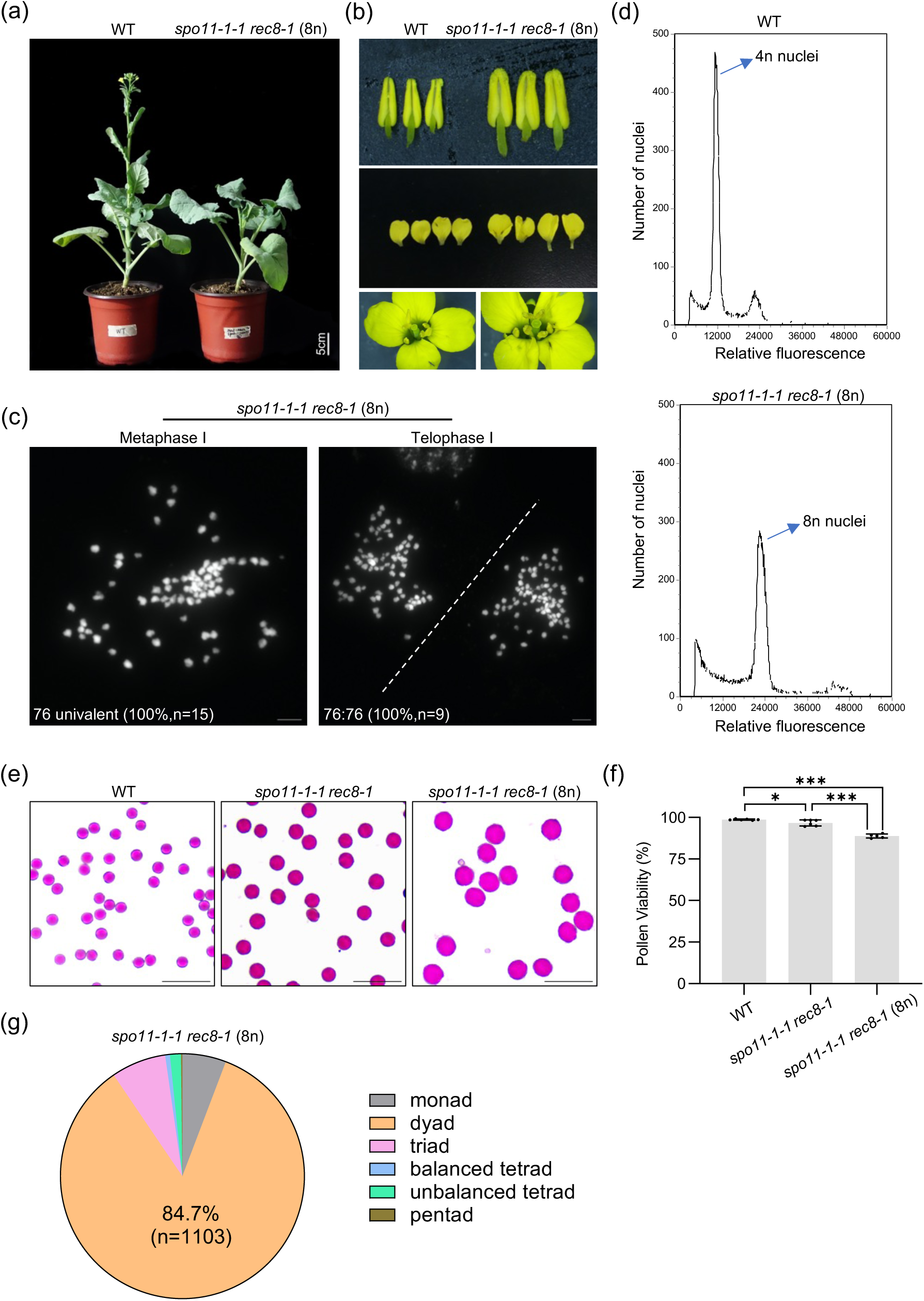
Phenotypic analysis of the progenies of *spo11-1-1 rec8-1* mutants. (a) The vegetative growth of WT and the progeny of *spo11-1-1 rec8-1* mutants. Since the ploidy of the progenies of *spo11-1-1 rec8-1* mutants are duplicated, they are referred to as *spo11-1 rec8-1* (8n). Bar: 5cm. (b) The floral organs of WT and *spo11-1-1 rec8-1* (8n) mutant plants. (c) Chromosome spreads of male meiosis in *spo11-1 rec8-1* (8n) mutants at metaphase Ⅰ and telophase Ⅰ. Bars: 5µm. (d) Flow cytometry analysis of WT and progenies of *spo11-1-1 rec8-1* mutants. (e) Pollen staining of WT, *spo11-1 rec8-1* and *spo11-1 rec8-1* (8X) mutant plants. Red pollen grains are fertile and grey pollen grains are aborted. Bars: 100µm. (f) Pollen viability of WT, *spo11-1-1 rec8-1* and *spo11-1-1 rec8-1* (8n) mutant plants. At least 6000 pollen grains were counted for each mutant. Asterisks indicate significant difference (Tukey’s multiple comparisons test, * *p*<0.05, *** *p*<0.001). (g) Pie chart showing the proportion of monad, dyad, triad, balanced tetrad, unbalanced tetrad, and pentad in *spo11-1-1 rec8-1* (8n) mutant plants.

**Figure S7.**
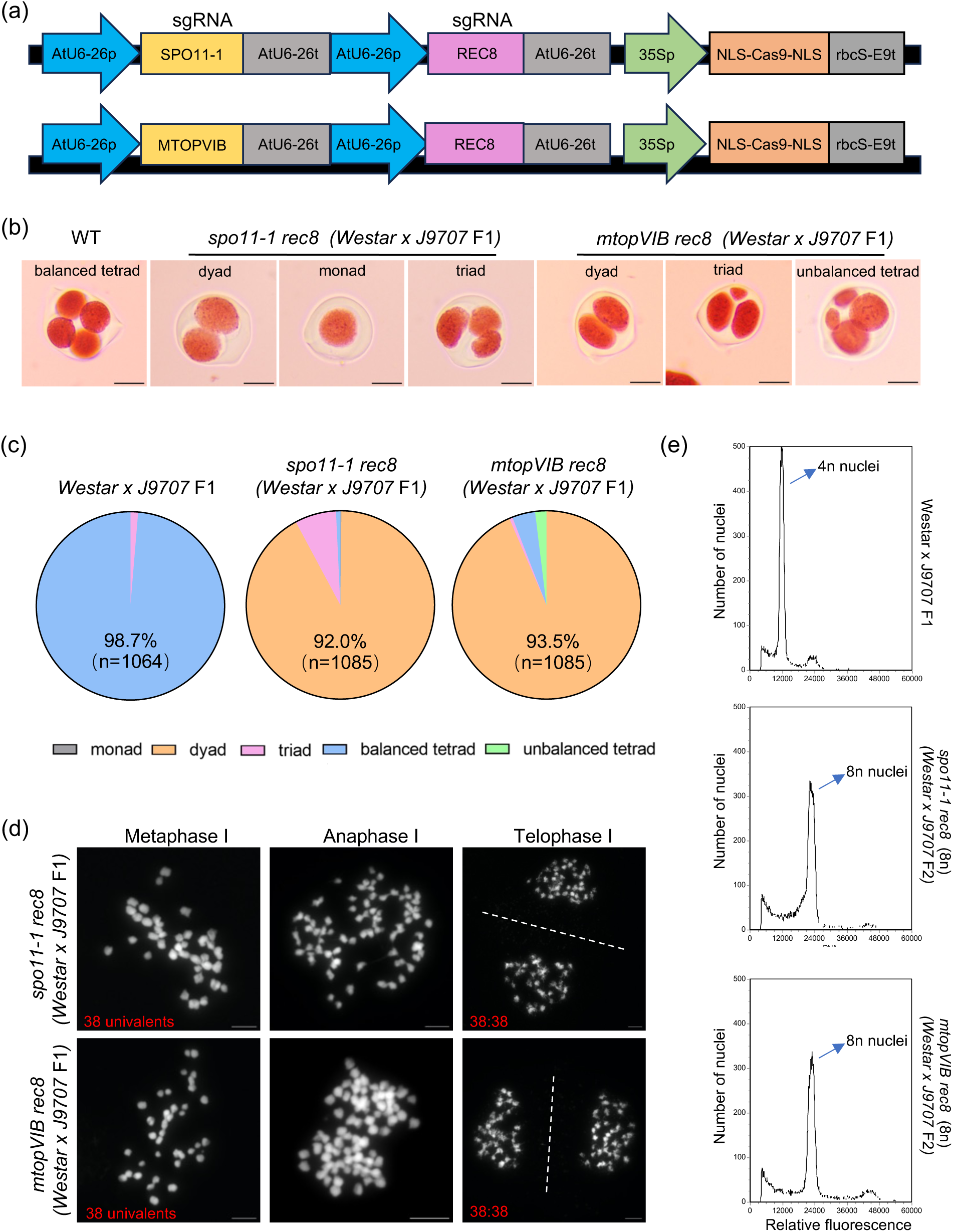
Generation of *MiMe* in F_1_ hybrids of two rapeseed cultivars *(Westar × J9707)*. (a) The structures of CRISPR-Cas9 vectors that simultaneously target *SPO11-1* or *MTOPVIB* and *REC8*. sgRNA, single guide RNA. (b) Male meiotic products at tetrad stage in the *Westar × J9707* F1 hybrids of WT, *spo11-1 rec8 (Westar × J9707)* and *mtopvib rec8 (Westar × J9707)* mutants. Bars: 5µm. (c) Pie charts showing the proportion of monad, dyad, triad, balanced tetrad, and unbalanced tetrad in the *Westar × J9707* F1 hybrids of WT, *spo11-1 rec8 (Westar × J9707)* and *mtopvib rec8 (Westar × J9707)* mutant plants. (d) Chromosome spreads of male meiosis in *spo11-1 rec8 (Westar x J9707)* and *mtopvib rec8 (Westar × J9707)* mutants in metaphase Ⅰ, anaphase Ⅰ, and telophase Ⅰ stages. Bars: 5µm. (e) Flow cytometry analysis of *Westar × J9707* F1, *spo11-1 rec8 (Westar x J9707* F2), and *mtopVIB rec8 (Westar x J9707* F2) mutant plants.

**Figure S8.**
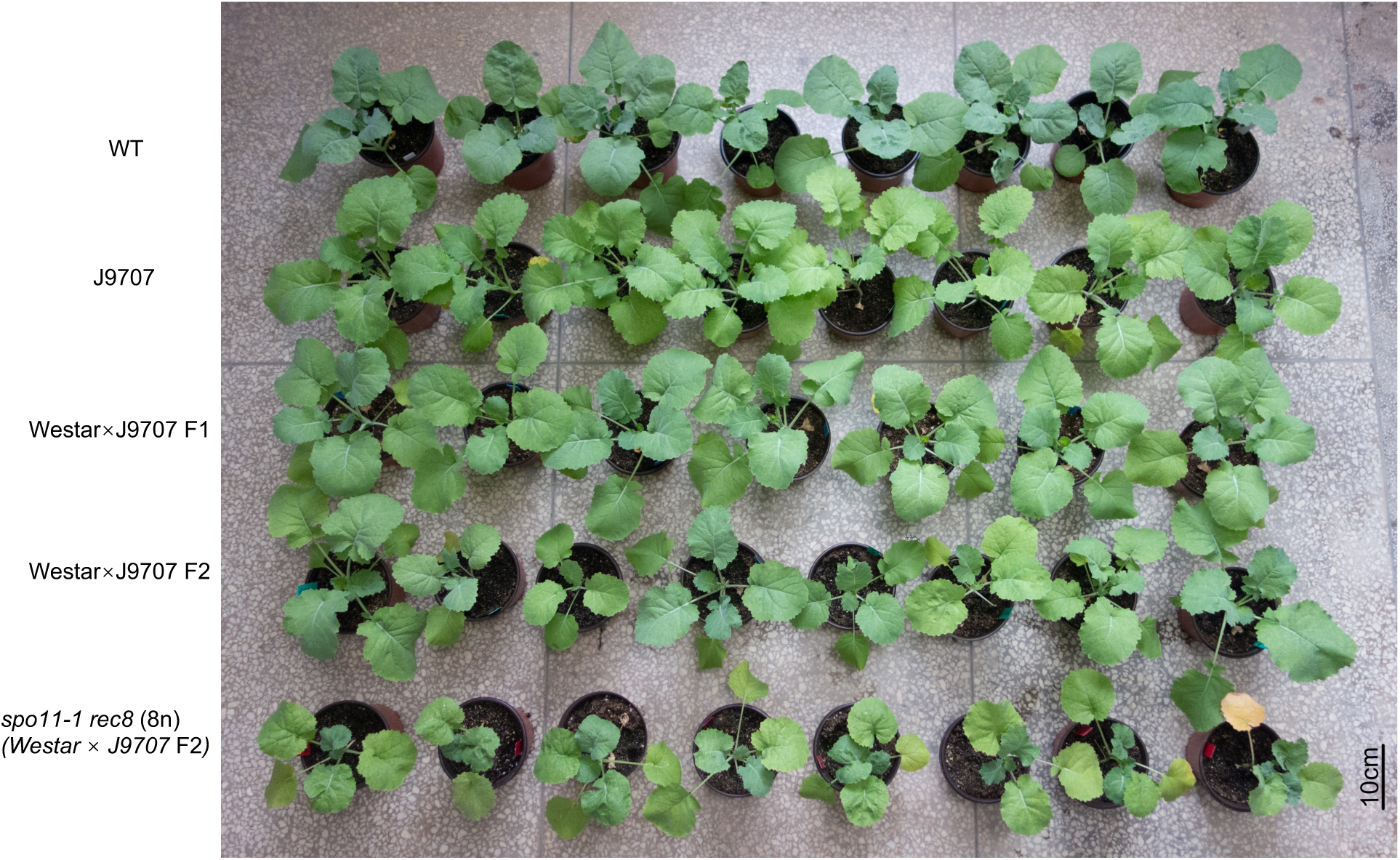
Growth comparisons of *westar*, *J9707*, *westar x J9707* F1, *westar x J9707* F2, and *spo11-1 rec8* (*Westar × J9707* F2) (8n) at 30 days after germination.

**Figure S9.**
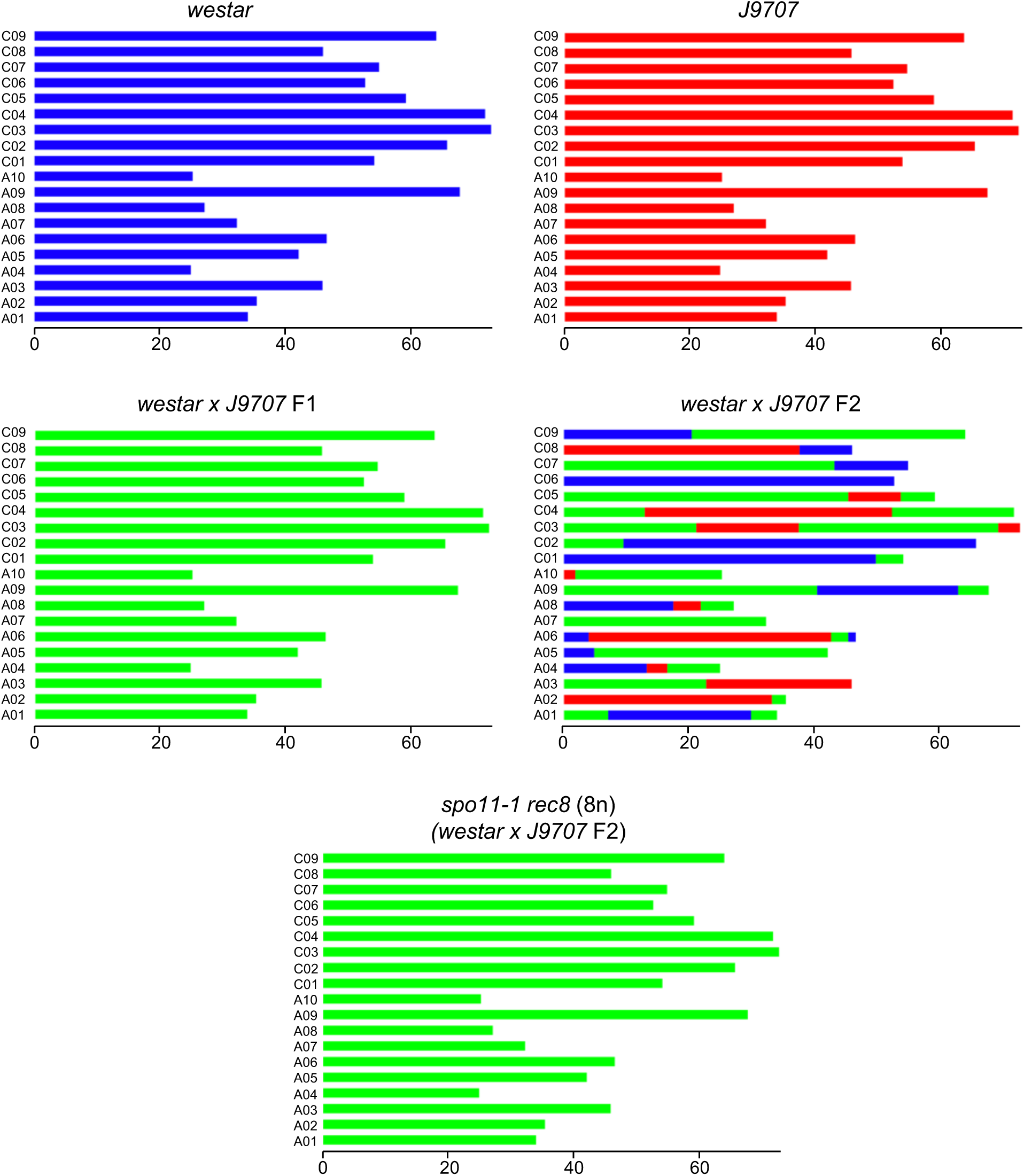
Whole-genome resequencing analysis for the genome-wide genotype landscape pf the hybrid *Brassica napus Westar x J9707* F1 (n=3), *Westar x J9707* F2 (n=3), *spo11-1 rec8* (*Westar x J9707* F2) (n=6), and their parents *Westar* (n=3) and *J9707* (n=3) plants. One example for each genotype is shown. The *Westar* alleles are indicated in blue, *J9707* in red, and *Westar x J9707* heterozygous alleles in green.

**Supplemental table 1.**
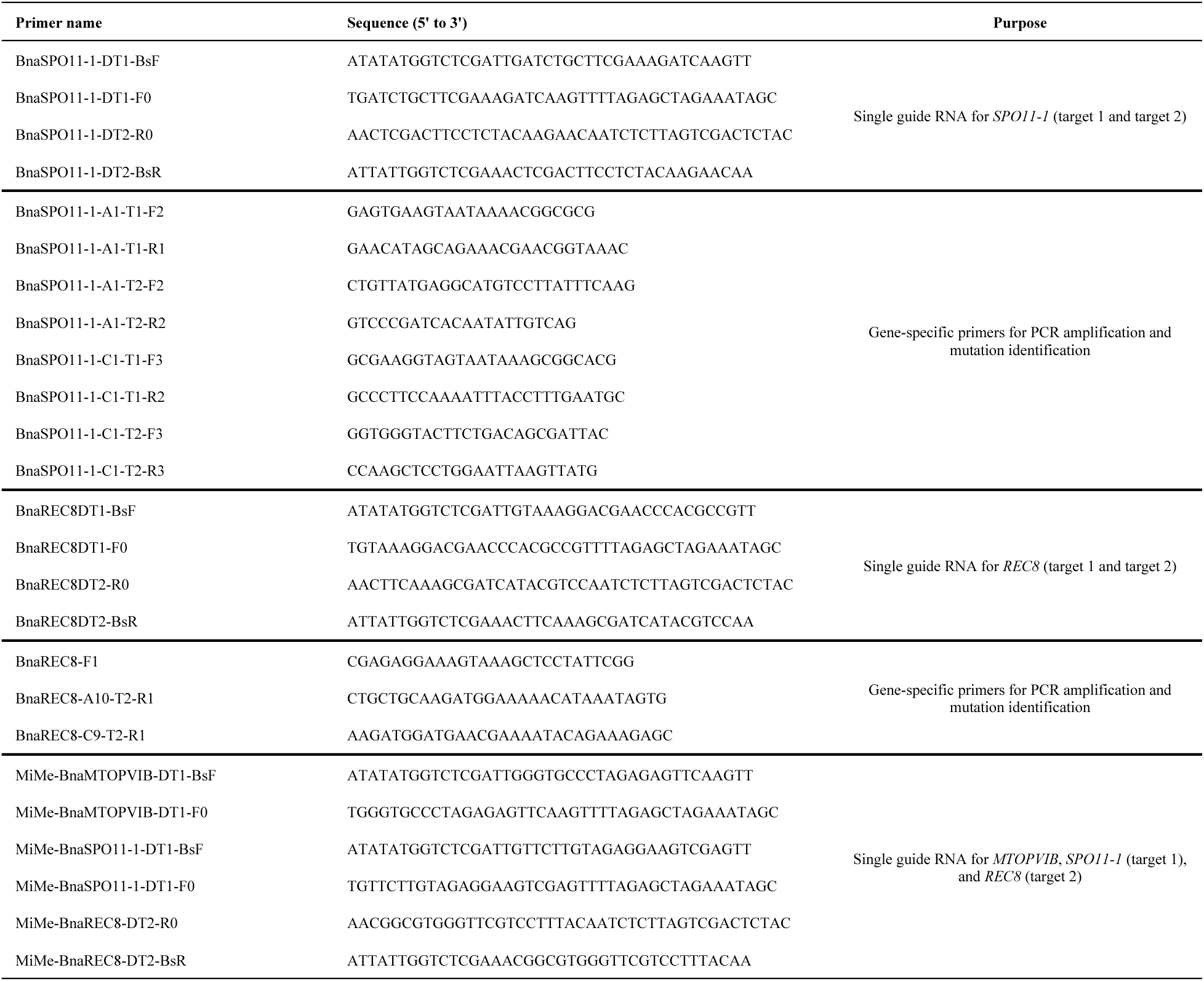
Primers used in this study.

